# LiP-Quant, an automated chemoproteomic approach to identify drug targets in complex proteomes

**DOI:** 10.1101/860072

**Authors:** Ilaria Piazza, Nigel Beaton, Roland Bruderer, Thomas Knobloch, Crystel Barbisan, Isabella Siepe, Oliver Rinner, Natalie de Souza, Paola Picotti, Lukas Reiter

## Abstract

Chemoproteomics is a key technology to characterize the mode of action of drugs, as it directly identifies the protein targets of bioactive compounds and aids in developing optimized small-molecule compounds. Current unbiased approaches cannot directly pinpoint the interaction surfaces between ligands and protein targets. To address his limitation we have developed a new drug target deconvolution approach based on limited proteolysis coupled with mass spectrometry that works across species including human cells (LiP-Quant). LiP-Quant features an automated data analysis pipeline and peptide-level resolution for the identification of any small-molecule binding sites, Here we demonstrate drug target identification by LiP-Quant across compound classes, including compounds targeting kinases and phosphatases. We demonstrate that LiP-Quant estimates the half maximal effective concentration (EC50) of compound binding sites in whole cell lysates. LiP-Quant identifies targets of both selective and promiscuous drugs and correctly discriminates drug binding to homologous proteins. We finally show that the LiP-Quant technology identifies targets of a novel research compound of biotechnological interest.

## INTRODUCTION

Unraveling the mechanism of action and molecular target of small molecules remains a major challenge in drug development. Current strategies to address this are often laborious and indirect. Genetic screens are common, offering the advantage of probing all potential targets simultaneously but as they measure the sum of many effects, it is difficult to discriminate direct versus *off-target* drug effects^1-3^. Direct target identification often requires additional time intensive steps such as compound modification or candidate protein purification, which in turn requires previous target evidence and can introduce confounding effects. Thermal proteome profiling (TPP) is an unbiased approach that can identify compound binding events in a complex proteome^4-6^ but it relies exclusively on alterations of a target’s thermal stability, involves complex melt curve analyses, and provides no structural information such as the compound binding site. This structural information is essential to understand bioactive compound interactions and to help on the development of optimized drug leads. Thus, the unbiased identification of drug-protein binding sites remains a challenge, especially in the context of its application to complex cellular systems.

To overcome this challenge, we have developed a novel technology (LiP-Quant) that identify compound targets and off-targets in an unbiased manner and can also provide additional compound-protein interaction information such as binding affinity and binding sites^7-9^.

## RESULTS

We recently presented an effective method to map small molecule binding proteins and binding sites directly from whole cell lysates of microbial organisms (LiP-SMap)^10^. The approach uses limited proteolysis (LiP) with a non-sequence specific protease and quantitative proteomics analysis to detect differential proteolytic patterns produced upon small molecule binding. The analysis reveals peptide fragments that change abundance upon compound binding (LiP peptides), and allow the identification of potential protein targets without prior hypothesis. The position of LiP peptides within the protein structure provides evidence for the small molecule binding sites^10^. However, this approach was only appropriate for the study of simple microorganisms and significant improvements were required to allow its application to more complex eukaryotic proteomes (e.g. human). Here we further developed LiP-SMap to enable the systematic investigation of protein-small molecule interactions in mammalian cell systems. To evaluate its performance we chose to focus on the identification of drug targets, an application of particular breadth and interest.

Here we present a LiP workflow optimized for the analysis of drug targets within human proteomes, called **LiP-Quant**. In this pipeline, protein lysates are exposed to a dilution series of a minimum of 7 drug doses followed by limited protease cleavage with proteinase K. Proteolytic patterns of drug targets should be altered upon addition of the drug, at least at the binding site, and true target peptides should show a change in abundance that correlates with drug concentration (**Figure 1A**).

**FIGURE 1:**
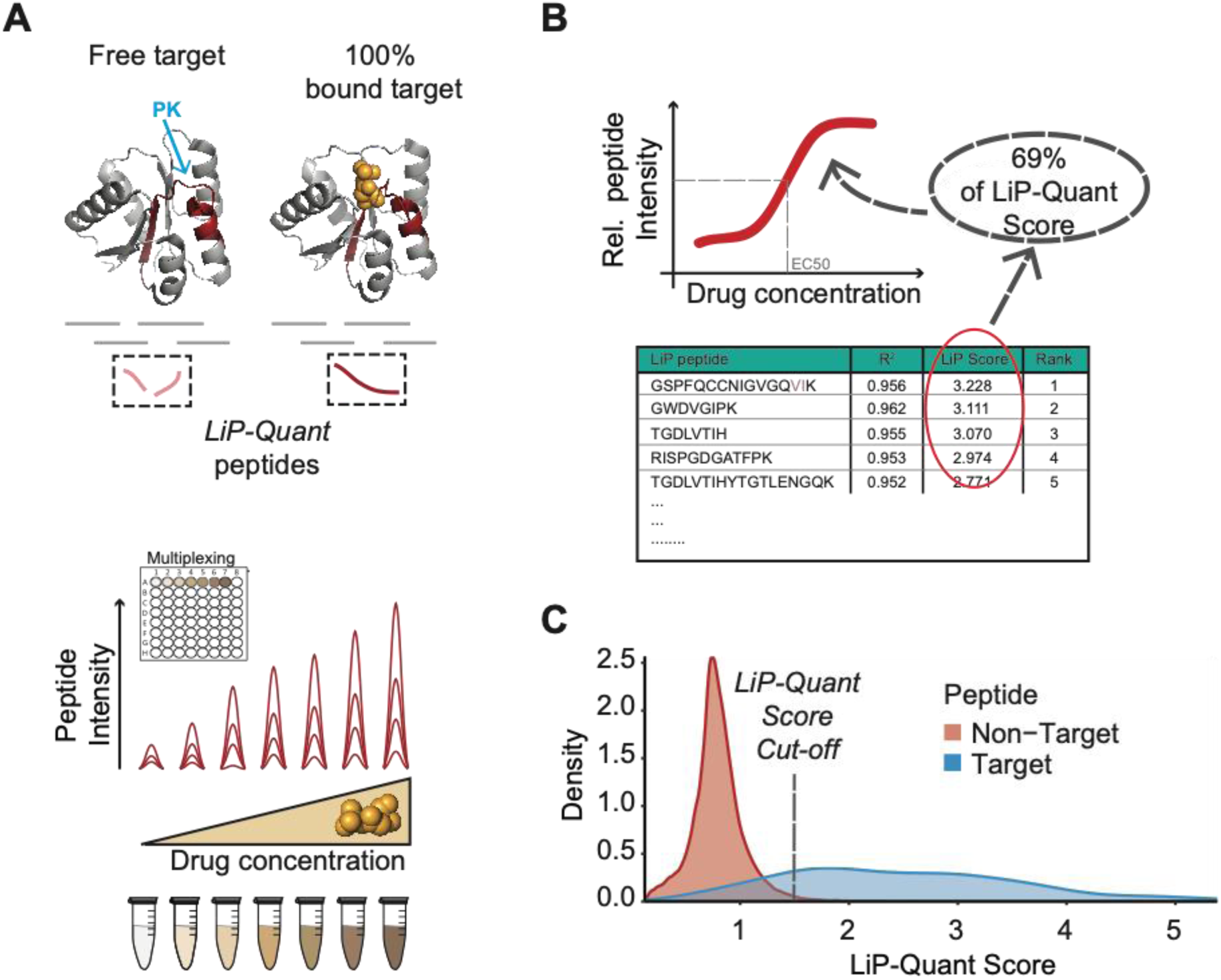
LiP-Quant, a platform for drug target identification. **1A:** Principle of LiP-Quant. Accessibility of proteinase K (PK) to structure specific cleavage sites in whole cell extracts changes depending on the fraction of drug molecules bound to target proteins, which increases with higher drug concentrations. This generates a population of LiP-Quant peptides at the PK cleavage sites that proportionally change abundance upon drug binding. Sample preparation for MS analysis follows a multiplexed workflow that is suitable for the processing of drug libraries. **1B:** Schematic depicting our scoring system (**LiP-Quant Score**) to rank LiP-Quant peptides. The predominant component (69%) of the LiP-Quant score is the correlation coefficient to a dose-response sigmoidal model. **1C:** Changes in peptide LiP-Quant score distribution for positive controls (drugs with known targets) enables the discrimination of target and non-target proteins. The distribution of LiP-Quant scores (Gaussian smoothed kernel density) from all HeLa experiments are shown, except that with the promiscuous binder staurosporine.

We have previously defined LiP peptides based on the statistical significance of their relative abundance changes upon metabolite binding^10^. However, that simplified approach limited its application for the investigation of relative simple microbial proteomes. With LiP-Quant we built a composite score (LiP-Quant score) based on machine learning of attributes that contribute to positive target identification. Several criteria (sub-scores) contribute to a peptide’s LiP-Quant score with the dominant component being correlation (R^2^) to a sigmoidal trend of the drug dose-response profile (69% of the LiP-Quant score) (**Figures 1B, S1A**). Sub-score weightings and an appropriate LiP-Quant score cutoff to distinguish target and non-target peptides were determined based on positive control experiments using drugs with known targets (**Figure S4A**) (see methods section). The combined LiP-Quant score enables direct comparison of LiP peptides with each other and allows more robust discrimination of genuine targets from random hits.

We observed that LiP-Quant scores show a bimodal distribution, with peptides from known target proteins clearly enriched in the high-scoring peak of the distribution (LiP-Quant score > 1.5), whereas non-target peptides are enriched in the low-scoring peak with a median of approximately 0.8 **(Figure 1C)**. Therefore, we defined putative LiP-Quant peptides as those with a LiP-Quant score above 1.5, which is the median score of non-target peptides plus three standard deviations. This threshold score ensures the presence of a minimal fraction of potential non-target peptides **(Table S1)** among candidates and a strong enrichment for genuine targets, with a Positive Predictive Value (PPV) of 31%, which is a three-fold increase in comparison to our previous LiP-SMap method (**Figure S1B**). Across our experiments, the PPV at the 1.5 LiP-Quant score threshold typically equates to that of the top 50 peptides (**Figure S1C**), yielding a ranked list of putative target peptides from the total of >100,000 peptides (5000 proteins) that were identified on average in each HeLa LiP-Quant experiment. By ranking peptides by LiP-Quant score, we prioritize peptides that are more likely to pinpoint the right target. The Top 10 peptides in the rank typically include 70% of true positives and the Top 100 only 18% (**Figure S1C)**.

We tested the macrolide immunosuppressive compound rapamycin with LiP-Quant, since the mechanism of binding to its known direct target (FK506-binding protein 1)^11,12^ is conserved across species. We quantified 30,209 peptides and 2618 proteins in *S. cerevisiae*, and 110,668 peptides and 5318 proteins in HeLa cells. Multiple LiP-Quant peptides, including the 5 top-scoring peptides, mapped to the known target of rapamycin in both *S.cerevisiae* (FRP1) (**Table S2**) and HeLa cells (FKBP1A) (**Figure 2A, 2B, S2A and S2B**) (**Table S1**), showing the ability of LiP-Quant to identify drug targets and the equivalence of the approach in yeast and humans.

**FIGURE 2:**
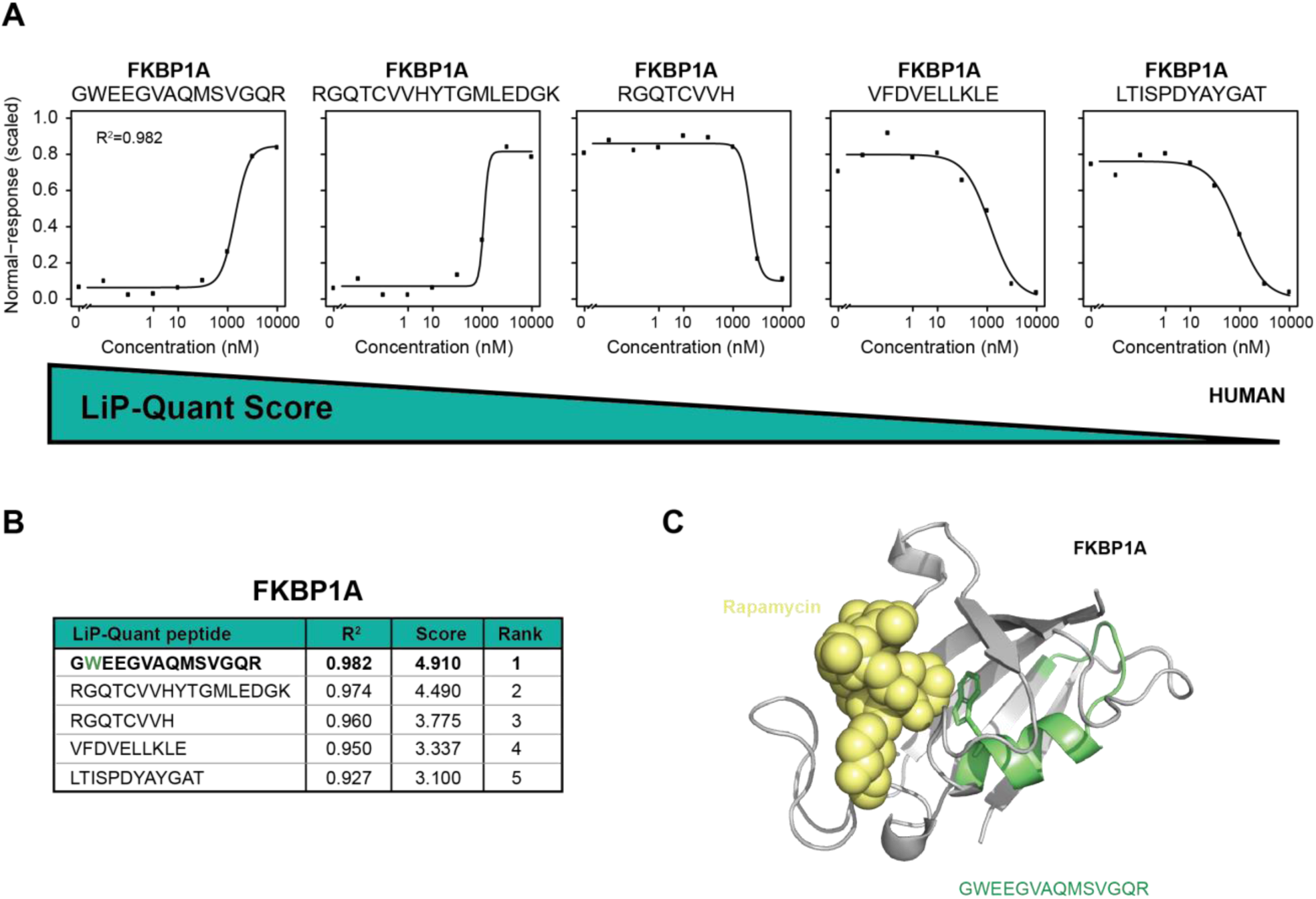
Benchmarking LiP-Quant in human cells. **2A:** Dose-response curves showing relative intensities of LiP-Quant peptides LiP-Quant peptides after partial proteolytic digestion of aliquots of HeLa lysates over a rapamycin concentration range. Curves of the top 5 LiP-Quant peptides ranked by LiP-Quant score, all of which are from the expected direct target FKBP1A, are shown. **2B:** LiP-Quant peptides ranking in positions 1-5 of the LiP-Quant experiment done with rapamycin in human cells. All 5 are FKBP1A peptides. **2C:** Structural model of the holocomplex of FKBP1A (PDBid: 2dg3) with rapamycin (yellow), showing the top-ranking peptide GWEEGVAQMSVGQR (green) from LiP-Quant experiments with rapamycin and FK506 (the top ranking peptide is the same). The minimal distance between the ligand and the top LiP-Quant peptide is 3.5 A.

The two top-scoring LiP-Quant peptides in each experiment, GWEEGVAQMSVGQR (FKPB1A, human) and GSPFQCCNIGVGQVIK (FRP1, yeast) are positioned in the known binding sites of rapamycin. The tryptophan residue of the top ranking FKBP1A LiP-Quant peptide **(Figure 2C)** and two isoleucine and valine residues of the first ranking FRP1 LiP-Quant peptide **(Figure S2C)** are in very close proximity with the compound atoms.

Beyond the known target of rapamycin FRP1 in yeast, we identified ARI1 and SYEC with high LiP-Quant scores (>2.5), which have not been previously characterized as targets of this compound **(Figure S2D)**. They could represent alternative rapamycin binding proteins (off-target effects) or be proteins that undergo secondary structural effects upon activation of the TOR1 pathway in the cell lysate. In order to discriminate between these two cases, the same LiP-Quant experiment was performed with lysates of a strain carrying a mutation in the *tor1* gene and a deletion of the *fpr1* gene *(tor1-1 Δfpr1)*^13^. In this strain, TOR signal transduction is impaired, including deletion of the direct receptor of the drug (FRP1), thus direct and secondary structural changes due to FRP1 binding and pathway activation do not occur. In the *tor1-1 Δfpr1* proteome, only peptide GDLVITEESWNK of ARI1 was detected as a hit. From this, we conclude that ARI1 is likely to be a secondary target of rapamycin (**Figures S2E and S2F**) (**Table S2**).

After establishing an improved analysis pipeline, we benchmarked it by testing the macrolide immunosuppressant FK506 in HeLa cells, as FK506 is known to target the same protein and binding site (FKBP1A) as rapamycin (**Table S1**). Target identification was very consistent between the compounds, with specific LiP-Quant peptides having similar scores and showing similar trends (i.e. fold increase or decrease) (**Figure S2G**). As for rapamycin, the highest ranking LiP-Quant peptide from FKBP1A (GWEEGVAMSVGQR) maps in very close proximity to FK506 (< 3Å) (**Figure 2C, Table S3**), confirming that LiP-Quant can consistently identify a known drug binding pocket. We identified three additional putative targets of both FK506 (CALR, TM263 and TXRD1) and rapamycin in HeLa cells (**Table S1**), which could represent additional *off-target* proteins of the two drugs in mammalian cells. Further work would be required to characterize the nature of these potential off-target interactions. We concluded that the quantitative LiP-Quant approach, with its associated data analysis pipeline, is suitable for drug-target deconvolution experiments in human cells with complex proteomes.

Next, we focused on kinases and phosphatases as drug targets of particular pharmacological interest, given their frequent dysregulation in disease, particularly in cancer. One common issue with kinase inhibitors (KIs) is their variable selectivity, as some KI drugs have one or two kinase targets in the cell, while others target hundreds simultaneously, making it difficult to determine their specific mode of action^14^. We used LiP-Quant to characterize KIs by comparing staurosporine, the best-studied promiscuous kinase-inhibitor, which binds the ATP binding sites of many kinases^15,16^, and the KI selumetinib, which specifically inhibits the MAP kinases MAP2K1 and MAP2K2 in different human cell lines ^17^. Amongst the top 25 peptides by LiP-Quant score we identified 18 peptides that map to 13 protein kinases for staurosporine and exclusively detected peptides from MAP2K1/2 for selumetinib, indicating that LiP-Quant can recapitulate the known selectivity profiles of the two compounds (**Figures 3A, 3B**). Interestingly, the protein NQO2 scored highly for both selumetinib (LiP-Quant rank #2) (**Figures 3C, S3B**) and staurosporine (LiP-Quant rank #8 and #14), which suggests it is a common off-target (**Figure S3B**). Although this finding is novel for these compounds, NQO2 is a confirmed off-target of several other kinase inhibitors including TBBz, DMAT and imatinib (Gleevec)^15,18,19^.

**FIGURE 3:**
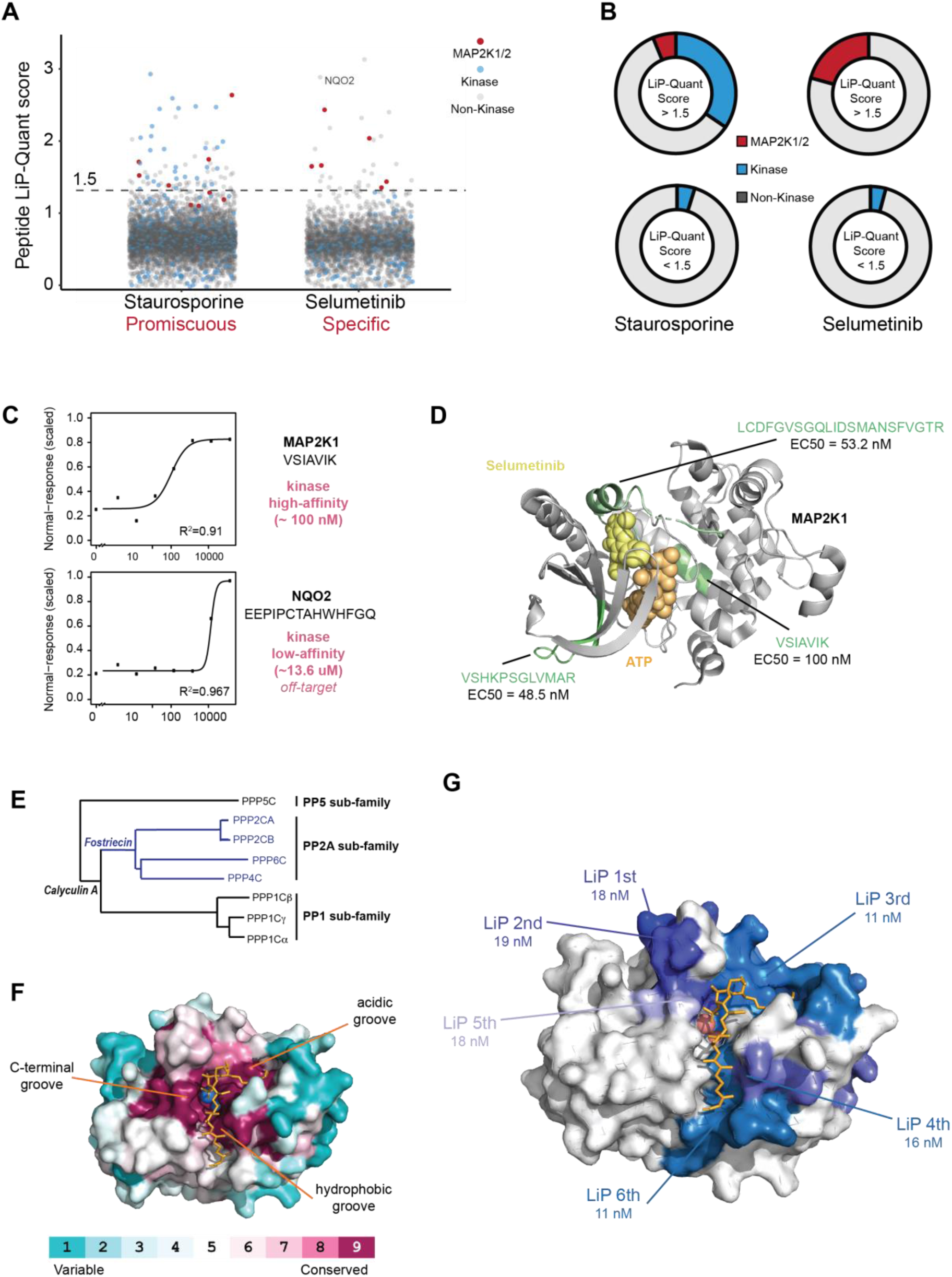
Probing druggable targets: kinases and phosphatases. **3A**: LiP-Quant captures druggable kinase targets with different specificity: Jitter-plots showing the distribution of LiP-Quant scores in LiP-Quant experiments done with the indicated kinase inhibitors. The blue dots show peptides assigned a Drug-score from any kinase, while gray dots show peptides assigned a LiP-Quant score from all classes of proteins. LiP-Quant peptides of MAPK proteins are shown in red. The LiP-Quant score cut-off for expected targets is shown with a dashed line. **3B:** Radar plots showing the proportion of LiP-Quant peptides corresponding to protein kinases among peptides with a LiP-Quant score higher or lower than the threshold score of 1.5. **3C**: Dose-response curves showing the relative intensities of the LiP-Quant peptides VSHKPSGLVMAR of mitogen-activated protein kinase kinase (MAP2K1) and EEPOPCTAHWHFGQ of NQO2 over a concentration range of selumenitib. The extrapolated EC50 for these two peptides are 100 nM and 13.6 μM respectively **3D:** Structural model of the holocomplex of MAP2K1 with the substrate ATP (orange spheres) and the drug selumetinib (yellow spheres) located in the active site and allosteric site of the kinase, respectively (PDBid: 4u7z) and its assigned LiP-Quant peptides. The LiP-Quant peptide LCDFGVSGQLIDSMANSFVGTR is present in both MAP2K1 and MAP2K2. **3E:** Phylogenetic tree (Clustal X neighbor joining tree) of the protein phosphatase (PP) family including subfamilies. The colors show the known sub-family selectivity of the compounds calyculin A and fostriecin. **3F**: Structural model of calyculin A bound to the PP1-gamma catalytic subunit (PDBID: 1it6). The surface has been colored in pink or blue according to amino acid conservation. calyculin A (represented with orange sticks) occupies the hydrophobic groove and the acidic groove on the molecular surface and adopts an extended conformation on the surface. **3G**: Surface representation of a structural model of calyculin A bound to PP2A with the surfaces corresponding to the top 6 ranking peptides by LiP-Quant score of PP2A (calyculin A experiment) colored with different tones of blue. Since the catalytic subunits of the PP family are highly structurally similar, the PP1-gamma subunit has been used as a model of the complex between PP2A and the drug after aligning the PP2A peptides to their homologs of PP1-gamma (pdbID: 1it6).

An unique advantage of the LIP based approach is its ability to identify interactions at peptide level resolution. We previously reported that the position of LiP peptides predicts the position of a binding site^10^. Here we confirm this observation as the position of the LiP-Quant peptides identified in the LiP-Quant experiments with staurosporine and selumetinib were in very close proximity to the known kinase inhibitors binding sites and also correctly pinpoint allosteric sites of MAP2K1, in the case of selumetinib (**Figures 3D and S3A**).

Next, we focused on inhibitors of serine/threonine phosphatases PP1 through PP6 (**Figure 3E**)^20^. The catalytic subunits of these enzymes are highly conserved at the active site region, which is elongated in a Y-shaped groove on the surface, thus lacking a defined binding pocket^21^ (**Figure 3F)**. We tested whether LiP-Quant could recapitulate the distinctive features of phosphatase inhibitors that bind only certain PP family members. Specifically, we tested calyculin A, which targets PP1 and PP2, and fostriecin, which binds PP2 and PP4 but not PP1^22,23^. As expected the top 15, and 21 of the top 25, peptides by LiP-Quant score from calyculin A all map to either PP1 or PP2A/B. The compound’s high target selectivity was also recapitulated, as no peptides from PP2C were found as hits (**Table S4**).

When we analyzed the structural location of the 6 highest ranking LiP-Quant peptides from PP2 for calyculin A, we found that they overlapped over the atypical extended Y-shaped groove of the PP2A active site. More specifically, they map over the acidic groove and the hydrophobic groove regions that correspond to the areas in direct proximity to calyculin A (**Figures 3F and 3G**). Similar results were obtained with fostriecin, as we observed the same LiP-Quant peptides or peptides mapping to the same region as calyculin A from PP2A/B (**Table S4**).

Finally, we investigated the ability of our new approach to provide binding affinity information. Since LiP-Quant yields a dose-response curve, it allows estimation of an EC50 value for a given compound (the concentration of drug at which we observe a variation of 50% of the maximum LiP signal). We asked whether these extracted EC50s matched the EC50 values previously reported in the literature. We used the kinase and phosphatase inhibitors assayed here (staurosporine, selumetinib, calyculin A and fostriecin) as test cases. The EC50 values we extracted for selumetinib for the top 3 peptides are 48.5, 53.2 and 100 nM, which are slightly above the EC50 of 41 nM measured with alternative methods (**Figure 3D**)^17^. Further, the inferred median EC50 values extracted from the top 5 peptides from PP2A/B and PP1 in the calyculin A LiP-Quant experiment are 18 nM and 63 nM, respectively and are approximately 10 fold higher to those measured *in vitro* (**Figure S3C)**^24^.

Although EC50 values estimated from LiP-Quant are not expected to be precise in absolute terms (see Discussion), our results show that they recapitulate very well the known relative affinities of calyculin A and fostriecin for the different PP family members (**Table S1**). The ratio between the calyculin A EC50 inferred by LiP-Quant for PP2A/B and PP1 closely reflects the previously reported 3.5-fold EC50 ratio difference between PP2A and PP1^24^. Moreover, we extrapolate a median EC50 (from the top 5 peptides) for PP2A/B of 20 nM for fostriecin, which is similar to both the published affinities of fostriecin for PP2A directly (0.2 to 40 nM) and to the relative affinity of calyculin A for PP2A (0.5 to 20-fold the EC50 of fostriecin)^25,26^. This ability of LiP-Quant to approximate absolute EC50 values and to effectively discriminate relative affinities between drug targets should uniquely help determine preferential target proteins of compounds.

In summary, we have shown that LiP-Quant assays are applicable to kinase inhibitors commonly used in cancer treatment. We correctly identify the targets of kinase inhibitors with different binding promiscuity, and predict possible off-targets as well as drug binding sites. LiP-Quant could differentiate highly homologous phosphatase proteins as targets of drugs with very subtle differences in specificity, illustrating the advantage of our peptide-centric approach beyond the mapping of binding sites.

Given its demonstrated abilities to identify and characterize drug targets, we envision a prominent role for LiP-Quant in the drug discovery pipeline. To evaluate this, we applied the approach to a research fungicide compound (BAYE-B004) that in phenotypic screens was found to inhibit *Botrytis cinerea* cell growth (Figure 4A), a mold that can devastate commercial fruit crops. Using LiP-Quant, we identified several proteins of interest as potential targets of BAYE-B004, including Bcin06g02870 (Figure 4B), predicted to be the B. cinerea homologue of casein kinase I. Only 7 peptides have a LiP-Quant score higher than 1.5. Among those, two LiP-Quant peptides from Bcin06g02870 have a low nanomolar EC50 of 6 and 5 nM (LiP-Quant rank #6 and #7), approximately 1000-fold lower than the EC50 of the rest of the LiP-Quant peptides above the 1.5 threshold (Table S5). We therefore tested further whether Bcin06g02870 was the primary target of the compound.

**FIGURE 4:**
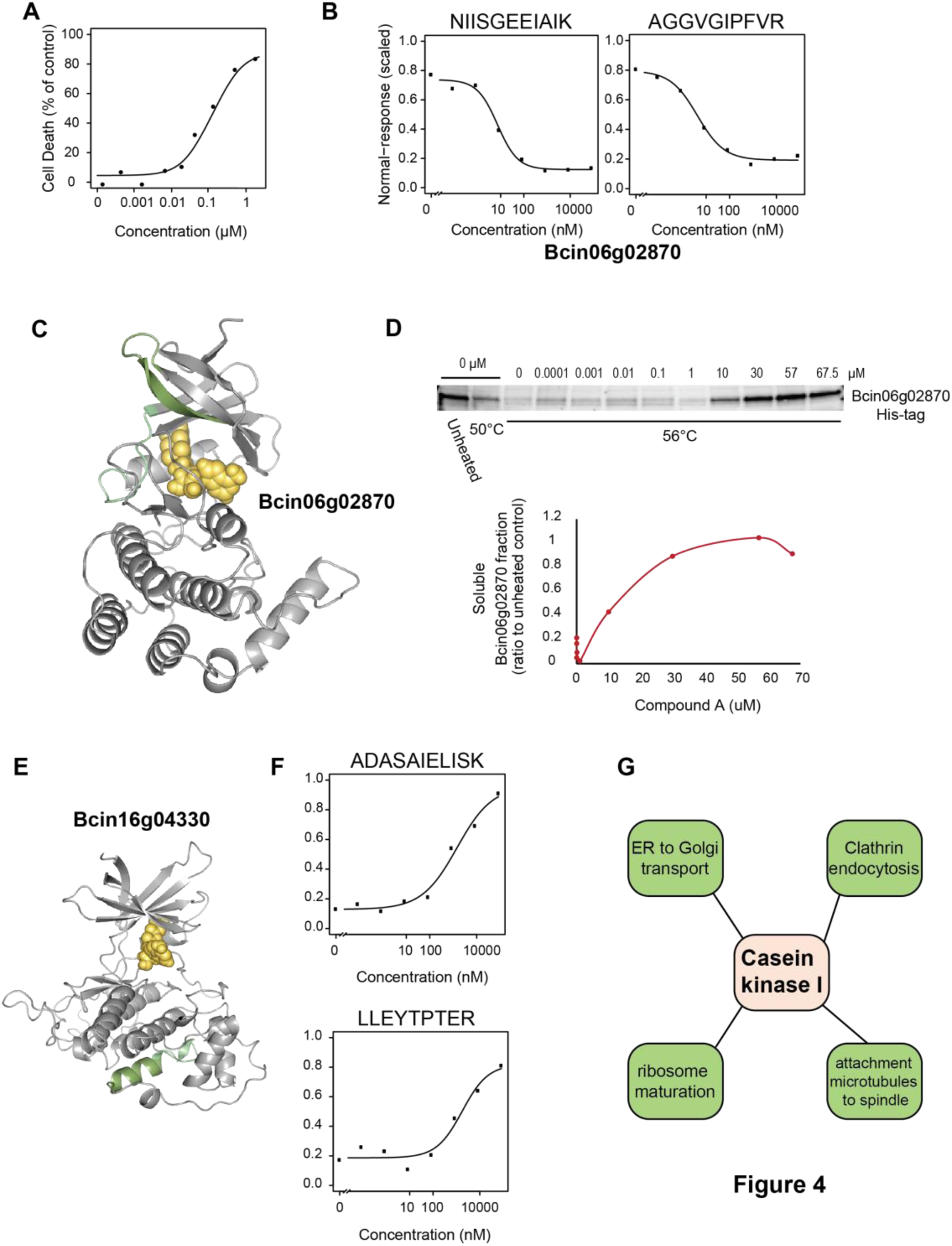
LiP-Quant of a novel fungicide. **4A:** Inhibition of growth in *Botrytis cinerea* cells upon treatment with increasing concentrations of a novel compound (BAYE-B004). **4B:** Dose-response curves showing relative intensities of the top two LiP-Quant peptides from Bcin06g02870, based on LiP-Quant score in the presence of increasing concentrations of a research fungicide. Bcin06g02870 is a serine/threonine kinase predicted to be casein kinase I. The extrapolated average EC50 for this protein is 6 nM. **4C:** The two top ranking LiP-Quant peptides (green) map directly to the predicted ATP-binding site (ATP in orange) of Bcin062870 predicted by homology modelling (template PDBid: 5cyz). **4D:** Thermal stability of His-tagged Bcin06g02870 upon treatment with increasing concentrations of BAYE-B004. Western blots and the corresponding quantification of the soluble fraction of Bcin06g02870 at 56°C are shown. **4E:** Structure of Bcin16g04330, a serine/threonine kinase, predicted by homology modelling (template PDBid: 4e7w), showing the position of the two top ranking LiP-Quant peptides for this protein (green) mapping outside the predicted ATP-binding site (ATP in orange). **4F:** Dose-response curves showing relative intensities of the top two ranking LiP-Quant peptides for Bcin16g04330 (see **Figure 4E**) The extrapolated average EC50 for this protein is 1.6 μM. **4G:** Proposed mechanisms of action by which inhibition of casein kinase I activity could inhibit cell growth.

Using a *Botrytis cinerea* cell line expressing a His-tagged version of Bcin06g02870, cellular thermal shift assays (CETSA)^5^ demonstrated that this protein is thermally stabilized upon treatment with BAYE-B004, confirming compound binding to Bcin06g02870 (**Figure 4D**). Based on structural modelling, both LiP-Quant peptides map very close to the predicted ATP-binding site of the predicted kinase (**Figure 4C**). Since this is a common binding site for kinase inhibitors, these data suggest that the compound is itself a kinase inhibitor (**Figures 3 and S3**). LiP-Quant also identified an additional kinase, Bcin16g04330, predicted to be the *B. cinerea* homologue of glycogen synthase kinase-3β (GSK-3b) (LiP-Quant rank #1 and #4, (**Table S5**) (**Figure 4E, 4F**). This kinase is likely not the primary target of BAYE-B004 as the extrapolated EC50 is several orders of magnitude higher at 1.6 μM (**Figure 4F**). Interestingly, the top 2 peptides of Bcin16g04330 map adjacent to each other but distal to the predicted ATP-binding site of the kinase, suggesting that the change in protease accessibility could be the result of either a compound binding-induced conformational change or a secondary binding or allosteric hindrance event (**Figure 4E**).

We propose that the observed inhibition in fungal cell growth upon BAYE-B004 treatment (**Figure 4A**) is due to mechanisms that are consistent with kinase inhibition, possibly targeting the kinases Bcin06g02870 (which is Casein kinase I in *Botrytis cinerea*) and Bcin16g04330 (GSK-3b). Casein kinase I has been shown to be required for cell viability in both budding yeast and cryptococcus neoformans^27,28^. Based upon these known functions we propose that inhibition of casein kinase I ultimately leads to reduced fungal cell growth (**Figure 4F**). Thus, by identifying putative targets of an uncharacterized drug, we demonstrate the relevance of LiP-Quant to drug development.

## DISCUSSION

Devising new unbiased chemoproteomic strategies that do not require chemical modifications of compounds and that simultaneously probe whole-proteomes will make the drug development pipeline more efficient. Here, we present LiP-Quant, a quantitative and unsupervised approach for drug target identification, based on the deconvolution of altered proteolytic patterns upon drug binding. In this approach, we identify drug targets via the automated evaluation of target peptide responses in terms of the quality of fit to the expected shape of a dose-response curve over a range of drug concentrations. We show that LiP-Quant substantially enriches for drug targets and can help to identify potential off-targets in both human cell lines and yeast.

LiP-Quant identifies peptides that undergo structural changes upon compound binding, identifying the drug binding sites for a benchmark set of protein kinase and phosphatase inhibitors. We could distinguish both the binding specificity and selectivity for the phosphatase inhibitors fostriecin and calyculin A, which interact with highly similar proteins that share a homologous binding site. This ability to accurately identify relative binding affinities for proteins, in particular among closely related protein families, is a highly beneficial feature when characterizing and refining drug leads.

We have found that LiP-Quant based EC50 calculations are typically consistent across different LiP-Quant peptides from a given target. However, EC50s estimated from LiP-Quant are often higher, by approximately an order of magnitude, than literature-reported values. EC50 values are generally measured in vitro with purified proteins, while those inferred from LiP-Quant are measured from lysates. The observed differences in EC50 values could thus simply be due to competition between different targets that occur in the lysate but not *in vitro*, or due to the effects of PTMs, protein-protein interactions, molecular crowding, binding of other small molecules or the presence of membranes. The EC50 estimated with LiP-Quant may in fact be a more physiological indicator of drug-target binding affinity, since protein lysates are a better model for the crowded environment of the cell than recombinant proteins *in vitro.*

We used LiP-Quant to discover the potential targets of a research fungicide, demonstrating the power of the approach when no prior target information is available. Our findings were corroborated by an additional target deconvolution approach and importantly, provide an explanation for the previously unknown mode of action of the drug. Interestingly, for the casein kinase-1 homologue, our approach suggests that the compound binds at the predicted ATP-binding pocket, a mechanism consistent with many well characterized kinase inhibitors. However, the top two peptides identified for another putative target, the glycogen synthase kinase-3β homologue, do not map to the predicted ATP-binding site. This illustrates that although binding site prediction is a powerful attribute of the method, such predictions should be orthogonally validated.

Collectively, this work demonstrates that LiP-Quant can be used to effectively identify protein drug-targets and characterize their binding properties across drug classes and species. These capabilities make LiP-Quant a powerful target deconvolution strategy with the potential to become an essential part of the drug development pipeline.

## METHODS

### Experimental model and subject details

#### Saccharomyces cerevisiae

cells were grown at 30°C in YPD media to early log phase from a single colony picked from a fresh YPD plate. Cells were harvested by centrifugation and carefully washed three times with ice-cold lysis buffer (100 mM HEPES pH 7.5, 150 mM KCl, 1 mM MgCl_2_). Cell pellets were resuspended in lysis buffer, and cell suspensions were extruded from a gauge needle to produce drops that were immediately flash frozen in liquid nitrogen.

#### Botrytis cinerea

(clone BO47), both wild-type and CK1 His-tagged, cells were cultured in potato dextrose agar (39 g/L, Oxoid #CM0139) at 21°C for 10 days. After 10 days growth the cells were suspended in 10 ml of GYPm liquid media (14.6 g/l D(+)-glucose monohydrate (VWR #24370.320), yeast extract (Merck #1.03753.0500), mycological peptone (Oxoid #LP0040)) and filtered (100 μm, Corning cell strainer) to harvest spores (final solution of 5 ×10^6^ spores/ml). This liquid culture was incubated for 24h at 110 rpm at 21°C. Cell mycelium was pelleted by centrifugation (5 min, 16,000 g, 21°C), media was removed and the pellet snap frozen in liquid nitrogen.

### Whole-proteome preparation for MS analysis

#### Saccharomyces cerevisiae

Liquid-nitrogen frozen beads of cell suspensions in lysis buffer (100 mM HEPES pH 7.5, 150 mM KCl, 1 mM MgCl_2_) were mechanically ground in cryogenic conditions with a Freezer Mill (SPEX SamplePrep 6875). Cell debris was removed by centrifugation (10 min, 20,000 g, 4°C). The sample preparation procedure was performed at 4°C.

#### HeLa and *Botrytis cinerea* cells

All biological samples were kept on ice through sample preparation. HeLa cell pellets (5 × 10^7^ cells) and *Botrytis cinerea* mycelium (3 × 10^7^ cells) were resuspended in 800 μl LiP buffer (100mM HEPES pH 7.5, 150mM KCl, 1mM MgCl_2_) and lysed by passing completely through a BD Precision glide syringe needle (27G) ten times, followed by 20 minutes incubation on ice. Lysate was cleared by centrifugation (16,000g at 4°C) for 4 minutes. Supernatant was retained in a new Eppendorf tube and the pellet was resuspended in 400 μl of LiP buffer for repeated lysis under the aforementioned conditions, including incubation and centrifugation. After centrifugation, supernatants were combined and protein amount was determined using a Pierce BCA Protein Assay Kit (cat #23225) according to manufacturer’s instructions.

### Limited Proteolysis under native conditions for global analysis of small molecule binding events

#### Saccharomyces cerevisiae

Cell lysates from at least three independent biological replicates were aliquoted in equivalent volumes containing 100 ug of proteome sample and incubated for 10 min at 25°C with the drug of interest. Proteinase K from Tritirachium album (Sigma Aldrich) was added simultaneously to all the proteome-metabolite samples with the aid of a multichannel pipette, at a proteinase K:substrate mass ratio of 1:100, and incubated at 25°C for 4 min. Digestion reactions were stopped by heating samples for 5 min at 98°C in a thermocycler followed by addition of sodium deoxycholate (Sigma Aldrich) to a final concentration of 5%. Samples were then heated again at 98°C for 3 min in a thermocycler. These samples were then subjected to complete digestion in denaturing conditions as described below.

#### HeLa and *Botrytis cinerea* cells

100 μg of protein lysate was aliquoted from a lysate pool for each of four independent replicates and incubated at room temperature (RT) with the compound of interest for 10 minutes. Proteinase K (1:100 ratio of enzyme to protein) was added and samples were incubated for a further 4 minutes. Samples were transferred to a heat block at 98°C for 1 minute, at which time proteinase K activity was quenched with an equal volume of 10% deoxycholate (to a final concentration of 5%) and incubated for a further 15 minutes at 98°C.

### Proteome preparation in denaturing conditions

Samples were removed from heat and reduced for 1 hour at 37°C with 5mM tris(2-carboxyethyl)phosphine hydrochloride followed by a 30 minute incubation at RT in the dark with 20 mM iodoacetamide. Subsequently, samples were diluted in 2 volumes of 0.1M ammonium bicarbonate (final pH of 8) and digested for 2 hours at 37°C with lysyl endopeptidase (1:100 enzyme: substrate ratio). Samples were further digested for 16 hours at 37°C with trypsin (1:100 enzyme: substrate ratio). Deoxycholate was precipitated by addition of formic acid to a final concentration of 1.5% and centrifuged at 16,000 g for 10 minutes. After transferring the supernatant to a new Eppendorf tube an equal volume of formic acid was added again and the centrifugation repeated. Digests were desalted using C18 MacroSpin columns (The Nest Group), or Sep-Pak C18 cartridges or into 96-well elution plates (Waters), following the manufacturer’s instructions and after drying resuspended in 1% acetonitrile (ACN) and 0.1% formic acid. The iRT kit (Biognosys AG, Schlieren, Switzerland) was added to all samples according to the manufacturer’s instructions.

### High pH reversed phase fractionation

Equal amounts of peptides were taken and pooled from the final LiP reaction digests for each treatment (e.g. 7 μg from each replicate for each condition), resulting in approximately 200 μg of total digest. This digest pool was fractionated into 10-12 fractions using high pH reversed phase chromatography with a Dionex Ultimate 3000 HPLC (Thermo Fisher, Waltham, United States) and an ACQUITY UPLC CSH C18 column (1.7 μm x 150 mm) from Waters (Milford, United States). In brief, a 25% ammonium hydroxide solution was used to adjust the pH of the digest pool to 10. The lysate was run on a 30-minute non-linear gradient, increasing from 1 to 40% ACN, at a flow rate of 0.3 ml per minute and a micro-fraction size of 30 seconds. After drying the individual fractions were resuspended in 1% ACN and 0.1% formic acid and Biognosys’ iRT kit was added.

### Mass spectrometric acquisition

#### Samples generated from HeLa and *Botrytis cinerea* cells

For DIA (Data Independent Acquisition) runs, 2 μg of LiP reaction digest from each sample was analyzed using an in-house analytical column (75 μm x 50cm). PicoFrit PicoTip Emitters (SELF/P Tip 10 μm) were packed with ReproSil-Pur C18-AQ 1.9 μm phase (Dr. Maisch, Ammerbuch-Entringen Germany) and connected to an Easy-nLC 1200. All experiments were run on a Q-Exactive HF mass spectrometer (Thermo Scientific) with the exception of the calyculin A data set, which was acquired on a Q-Exactive HF-X. Peptides were separated by a 2-hour segmented gradient at a flow rate of 250 nl/min with increasing solvent B (0.1% formic acid, 85% ACN) mixed into solvent A (0.1% formic acid, 1% ACN). Solvent B concentration was increased from 1% after 3 minutes according to the following gradient: 4% over 3 minutes, 5% for 3 minutes, 7% for 4 minutes, 9% for 5 minutes, 11% for 8 minutes, 16% for 19 minutes, 26% for 41 minutes, 29% for 9 minutes, 31% for 6 minutes, 33% for 5 minutes, 35% for 4 minutes, 38% for 4 minutes, 40% for 3 minutes, 44% for 3 minutes, 55% for 3 minutes and 90% in 10 seconds. This final concentration was held for 10 minutes followed by a rapid decrease to 1% over 10 seconds, which was then held for 5 minutes to finish the gradient. A full scan was acquired between 350 and 1650 m/z at a resolution of 120,000 (ACG target of 3e6 or 7 ms maximal injection time). A total of 37 DIA segments on HF were acquired at a resolution of 30,000 (ACG target of 3e6 or 47 ms maximal injection time) and 42 on the HF-X (ACG target of 3e6 or 55 ms maximal injection time). The normalized collision energy was stepped at 25.5, 27, 30. First mass was fixed at 200m/z.

For DDA (Data Dependent Acquisition) runs, peptides were separated by the same 2-hour segmented gradient as utilized above for DIA runs with the exception that the final 1% solvent B flow was held for 4 minutes and 40 seconds (rather than 5 minutes). All experiments were run on a Q-Exactive HF mass spectrometer (Thermo Scientific) with the exception of the rapamycin (Q-Exactive HF-X) and FK506 data sets (Q-Exactive). A top 15 method was used across a scan range of 350 to 1650 m/z with a full MS resolution of 60,000 (ACG target of 3e6 or 20 ms injection time). Dependent MS2 scans were performed with a resolution of 15,000 (ACG target of 2e6 or 25 ms injection time) with an isolation window of 1.6 m/z and a fixed first mass of 120 m/z.

Peptide samples generated from *Saccharomyces cerevisiae* were analyzed on an Orbitrap Q Exactive Plus mass spectrometer (Thermo Fisher Scientific) equipped with a nano-electrospray ion source and a nano-flow LC system (Easy-nLC 1000, Thermo Fisher Scientific). MS data acquisition in DDA and DIA modes was essentially carried out as in Piazza et. al. 2018.

### Mass spectrometric data analysis

DIA spectra were analyzed with Spectronaut X (Biognosys AG)^29^ using the default settings. In brief, retention time prediction type was set to dynamic iRT (adapted variable iRT extraction width for varying iRT precision during the gradient) and correction factor for window 1. Mass calibration was set to local mass calibration. The false discovery rate (FDR) was estimated with the mProphet approach^30^ and set to 1% at both the peptide precursor and protein level. Statistical comparisons were performed on the modified peptide level using fragment ions as quantitative input. The DDA spectra were analyzed with the SpectroMine (Biognosys AG) software using the default settings with the following alterations. Digestion enzyme specificity was set to Trypsin/P and semi-specific. Search criteria included carbamidomethylation of cysteine as a fixed modification, as well as oxidation of methionine and acetylation (protein N-terminus) as variable modifications. Up to 2 missed cleavages were allowed. The initial mass tolerance for the precursor was 4.5 ppm and for the fragment ions was 20 ppm. The DDA files were searched against the human UniProt fasta database (updated 2018-07-01) and the Biognosys’ iRT peptides fasta database (uploaded to the public repository). The libraries were generated using the library generation functionality of SpectroMine with default settings.

### Establishing criteria to be used for positive target identification in LiP-Quant

All HeLa data sets were first analyzed for differentially regulated peptides between the highest drug concentration and vehicle using Spectronaut’s statistical testing performed on the modified peptide sequence level using fragment ions as the smallest quantitative units. This candidate peptide list was filtered based upon q-value < 0.01 and an absolute log_2_ fold-change > 0.58. Each peptide in this filtered list was then subjected to dose response correlation testing (using the “drc” package (https://www.r-project.org)) on all peptides (modified sequence with fragments ions as quantitative units) at every drug concentration to establish a sigmoidal correlation coefficient.

As the ground truth (target proteins) was known for the drugs tested in HeLa lysates each protein identified in each data set was annotated as either a known target or non-target and from this a contaminant database, or LiP-protein frequency library (PFL), was built. To do so, the same statistically filtered list of differentially regulated peptides as above was used and proteins that were present but not specific for the drug being tested were quantified and assigned a PFL (contamination) score. For example, a protein that showed differential regulation in 9 of 11 ground truth experiments (several experiments were performed more than once) was assigned a contamination score of 9/11 or 81.8% (**Table S6**), proteins that never appeared as contaminants in any experiment were not included in the PFL-library. This library enabled the quantitative down-weighting of proteins that were frequently present in LiP experiments but not specific for the drug being tested. We observed high correlation between proteins identified as likely contaminants in the PFL of our LiP-Quant experiments (**Table S6**) and those previously identified as common contaminants in affinity purification mass spectrometry (such as chaperone and structural proteins) (Mellacheruvu et al., 2013).

To establish the criteria that contribute to the identification of drug targets, we split our dose response experimental data (filtered based on q-value and log_2_ fold-change and PFL annotated as mentioned above) into two independent data sets to train our classifier; training set A included the drugs calyculin A, rapamycin and staurosporine and training set B included FK506, selumetinib and fostriecin (**Figure S4A**). For each training set the data was combined and we used linear discriminant analysis (LDA) to build classifiers based upon all potential unique peptide/protein features (e.g. dose-response correlation, PFL frequency, protein coverage, etc). For each training set, known drug targets were selected as a positive training set, resulting in 95 modified sequences for training set A and 33 for training set B. We also randomly sampled 400 background modified sequences as a negative training set from each training set. The features were calculated and stabilized to a defined range between 0 and 1. The LDA-based machine learning was performed five times for each training data set with resampling of the negative training set each time. The identified criteria were consistent across all LDA analyses (**Figure S1A**) and the contribution weights for each of the features from the five LDA analyses was averaged. The relative contributions of each parameter to the LiP-Quant score was very stable across the training sets (**Figure S4B**). We termed the linear classifier the “LiP-Quant Score” in this study. The weights were adjusted such that the combined linear classifier could reach a maximum value of 6. These weightings were incorporated into the analysis pipeline (see below) and verified independently on the other positive control data sets (i.e. training set A was verified on the data sets comprising training set B and vice versa) (**Figure S4A**). LiP rankings using both training set analysis parameters were similar across all data sets (**Figure S4C**).

Using this approach, we established four classifiers that contribute to positive drug target identification (**Figure S1A**): (I) correlation of fit with a dose response binding model, (II) the presence of the identified protein in the LiP-protein frequency library, (III) the number of peptides from an identified protein showing regulation that are in the top ten percent of all peptides ranked by q-value in the Spectronaut filtered statistical test (see above) and (IV) the statistical significance (q-value) of the relative peptide abundances between drug and vehicle-treated samples. As training set A contained a larger positive training set (i.e. there were more known drug target peptides identified) the weightings calculated for this training set were used for all subsequent analyses.

### Automated peptide/protein ranking of LiP dose response experiments

Using the criteria and weightings established from our training data sets we wrote in-house scripts in R to calculate in an unbiased manner the individual peptide sub-scores for each LiP-Quant experiment. As these experiments contained on average over 100,000 peptides, peptides were first filtered based upon differential abundance from the Spectronaut statistical testing table (q-value < 0.01 and an absolute log_2_ fold-change > 0.46) using statistical comparisons against vehicle control for a range of drug concentrations (IC_50_ through 1000-fold the IC_50_, or the range closest to this). Each peptide in this narrowed down putative candidate list was then subjected to full LiP-Quant analysis using the four weighted criteria (**Figure S1A**) described above and a final LiP-Quant score for each peptide was calculated.

This final analysis pipeline enabled the selection and ranking of the most relevant peptides and proteins per experiment. The combined LiP-Quant score enables direct comparison of LiP peptides with each other and allows more robust discrimination of genuine targets from random hits. Ranking on the protein level was performed using the best LiP-score per protein, only. All half maximal effective/inhibitory concentrations (EC_50_/IC_50_) were calculated using the “drc” package (https://www.r-project.org).

### Criteria used for establishing a LiP threshold score

Aggregating results from five positive control experiments (rapamycin, calyculin A, selumentinib, FK506 and fostreicin) conducted in HeLa lysate and analyzed with our LiP-Quant pipeline, we found that LiP scores show a bimodal distribution. Staurosporine was excluded from the threshold calculation as it shows a level of promiscuity (binding potentially hundreds of kinases) that is rare among drugs, making it difficult to ascertain if low scoring peptides are genuine targets that were not detected or kinases that are not bound by the drug. As this difficulty in interpreting non-target peptides could bias the threshold calculation the data set was excluded. Peptides from known target proteins show a clear enrichment in the high-scoring peak of the distribution (LiP-Quant score > 1.5), whereas all other peptides are enriched in the low-scoring peak of the distribution with a median of approximately 0.8 (**Figure 1B**). We defined a threshold score of 1.5 by taking the median LiP-Quant score from the aforementioned experiments, plus three standard deviations, to ensure minimal (< 1%) non-target peptide presence (**Table S1**). Although the approach ensures a strong enrichment for genuine targets, it should be noted that some peptides from these targets are expected below a LiP-Quant score of 1.5 as both LiP-Quant and non-LiP-Quant peptides can be expected from genuine target proteins.

### Structural models

The amino acid conservation in the structural model of calyculin A bound to the PP1-gamma catalytic subunit has been calculated using the ConSurf algorithm (Landau 2005).

### Definition of Positive Predictive Value (PPV)

We defined the positive predictive value (PPV) as the ratio between the number of true positive peptides and the sum of false positives (FP) and true positives (TP) identified by LiP-Quant (TP/ (TP + FP)).

### Cellular thermal shift assay (CETSA)

*Botrytis cinerea* BO47 (CK1 His-Tagged) cell suspension was adjusted to 1 × 10^6^ sp/ml GYPm and incubated for 24h at 21°C (110 rpm). 12.5 × 10^6^ cells were treated with BAYE-B004 (at various concentrations from 0.0001 to 67.5 μM) or control (1% DMSO) for the final 20 minutes of the 24h growth period. Cells were harvested by filtration (100 μm) and rinsed with 15 ml of ice cold HEPES buffer (0.1 M HEPES, 50 mM NaCl, pH 7.5). Harvested mycelium was resuspended in 3.5 ml HEPES buffer and kept on ice. 500 μl of each concentration was transferred to a 2 ml Eppendorf tube and heated to 56°C on a thermoshaker for 3 minutes, an additional aliquot from each concentration was left unheated. After heating, cells were kept on ice for 3 minutes, snap frozen in liquid nitrogen, lyophilised overnight and then stored at -80°C until protein extraction.

Lyophilised mycelium was lysed using a Retsch mixer mill (MM 400) with 3 mm tungsten carbide beads (30 Hz for 3 seconds, two cycles), then 500 μl of cold protein extraction buffer (50 mM HEPES, 50 mM NaCl, 0.4% NP-40) was added. Lysate was incubated for 10 minutes at 25°C, centrifuged (10 minutes, 14,000g) and the supernatant was retained. The lysate was further centrifuged (20 minutes, 73,400g) to eliminate insoluble proteins. The supernatant was collected and protein concentration was determined using the Qubit protein assay kit (#Q33211) and stored at -20°C.

Target engagement was assessed by western blot. In brief, 17 μg of protein per treatment was loaded onto a TGX (4-20%) stain free gel (Bio-Rad, #4568094) and run at 250V for 25 minutes. Proteins were transferred to a nitrocellulose membrane using the Trans-Blot Turbo system according to the manufacturer’s instructions (Bio-Rad, # 1704271). The membrane was probed using a monoclonal anti-polyhistidine-peroxidase antibody (1:2000, clone HIS-1, Sigma, A7058). The membrane (target protein) and gel (loading control) were imaged using a ChemiDocXRS camera and quantified using ImageJ ^31^

### Cell viability (IC50) assay

*Botrytis cinerea* BO5.10 (2 × 10^3^ cells/ml) mycelium in GYPm liquid media (200 μl) was cultured at 21°C without shaking in a micro-titer plate. Optical density was measured at 620 nm (Tecan M1000 plate reader) at the beginning of the culture period (day 0) and immediately inoculated with 2 μl of BAYE-B004 to obtain final concentrations (μM) of 1.2234, 0.40745, 0.13582, 0.04527, 0.01509, 0.00168, 0.00056, 0.00019 and 0 respectively. The culture was grown for three days at 21°C after inoculation at which point the optical density was measured again. Inhibition of cell growth was calculated at each.

## MATERIALS

Frozen HeLa cell pellets were purchased from Ipracell (Belgium). All chemicals and compounds were purchased from Sigma-Aldrich unless specified otherwise. Pierce BCA Protein Assay Kit was purchased from Thermo Fisher Scientific. Lysyl endopeptidase was purchased from Wako Pure Chemical Industries. Selumetinib and staurosporine were purchased from Lubio Science and Cell Guidance systems respectively. Fostrecin was purchased from AdipoGen. BAYE-B004 was produced by Bayer Crop Science. Sequencing grade trypsin was purchased from Promega.

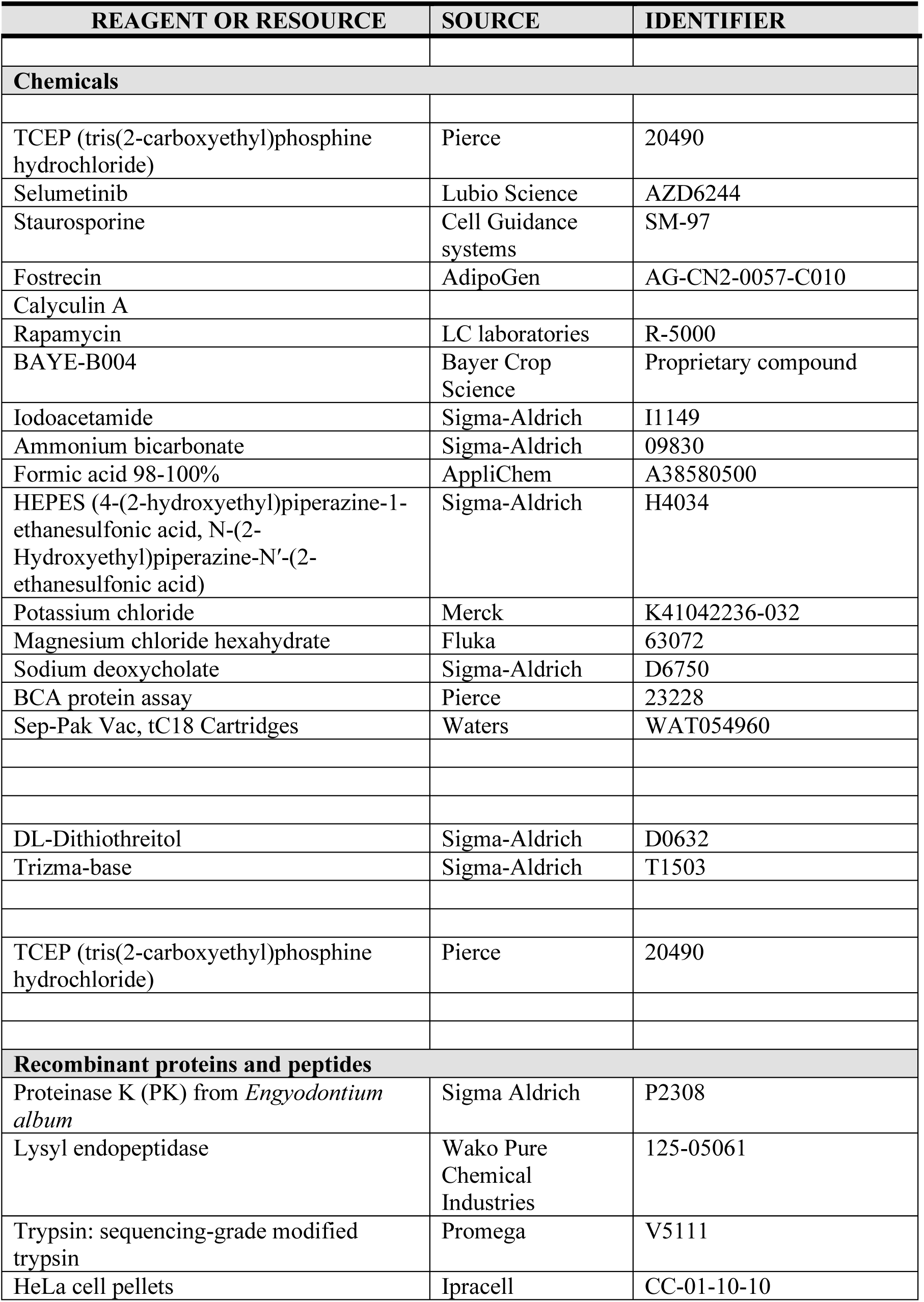
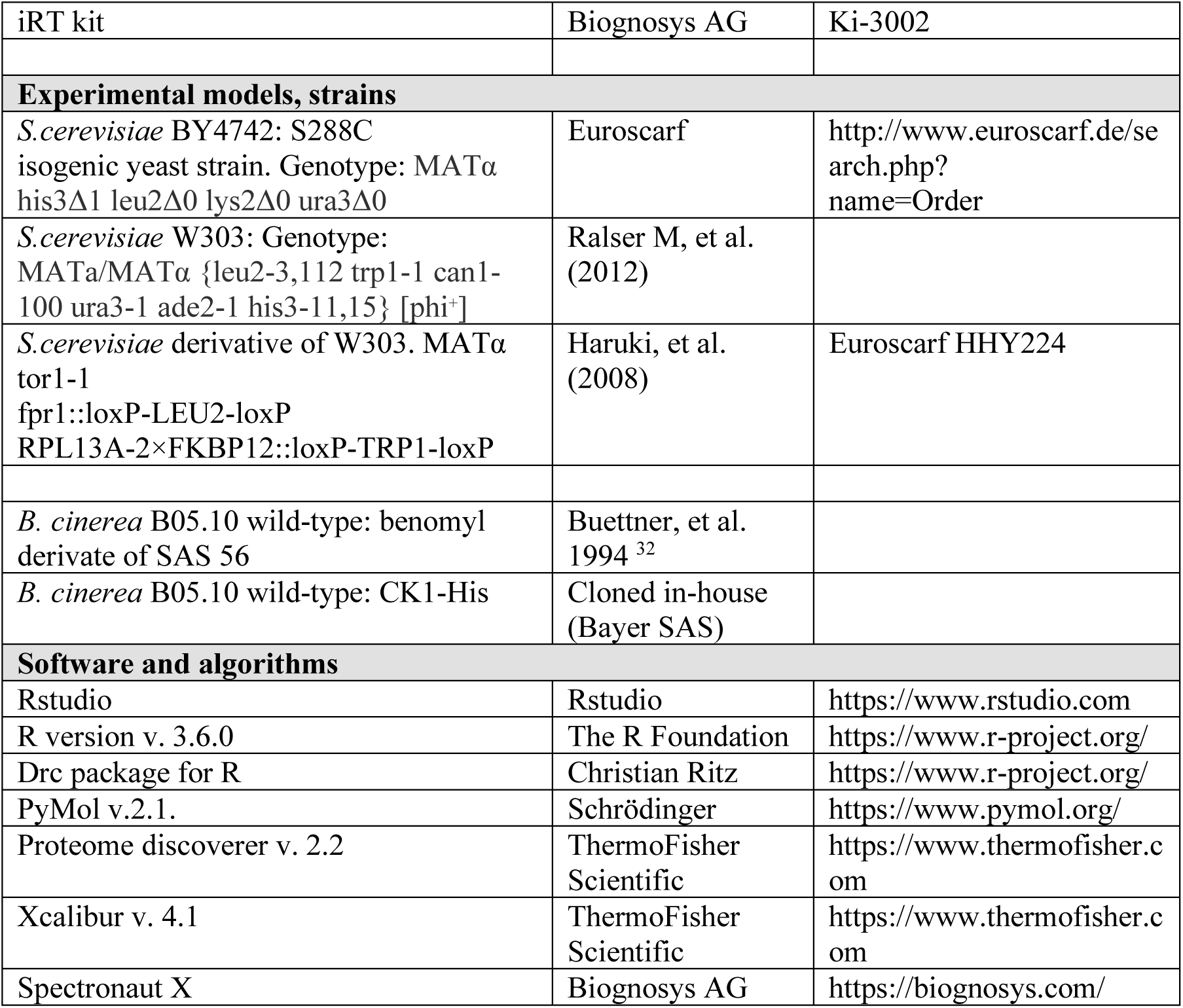

