## Supplementary Figures for "LiP-Quant, an automated chemoproteomic approach to identify drug targets in complex proteomes"

Chemoproteomics is a key technology to characterize the mode of action of drugs, as it directly identifies the protein targets of bioactive compounds and aids in developing optimized small-molecule compounds. Current unbiased approaches cannot directly pinpoint the interaction surfaces between ligands and protein targets. To address his limitation we have developed a new drug target deconvolution approach based on limited proteolysis coupled with mass spectrometry that works across species including human cells (LiP-Quant). LiP-Quant features an automated data analysis pipeline and peptide-level resolution for the identification of any small-molecule binding sites, Here we demonstrate drug target identification by LiP-Quant across compound classes, including compounds targeting kinases and phosphatases. We demonstrate that LiP-Quant estimates the half maximal effective concentration (EC50) of compound binding sites in whole cell lysates. LiP-Quant identifies targets of both selective and promiscuous drugs and correctly discriminates drug binding to homologous proteins. We finally show that the LiP-Quant technology identifies targets of a novel research compound of biotechnological interest.

### SUPPLEMENTARY FIGURES

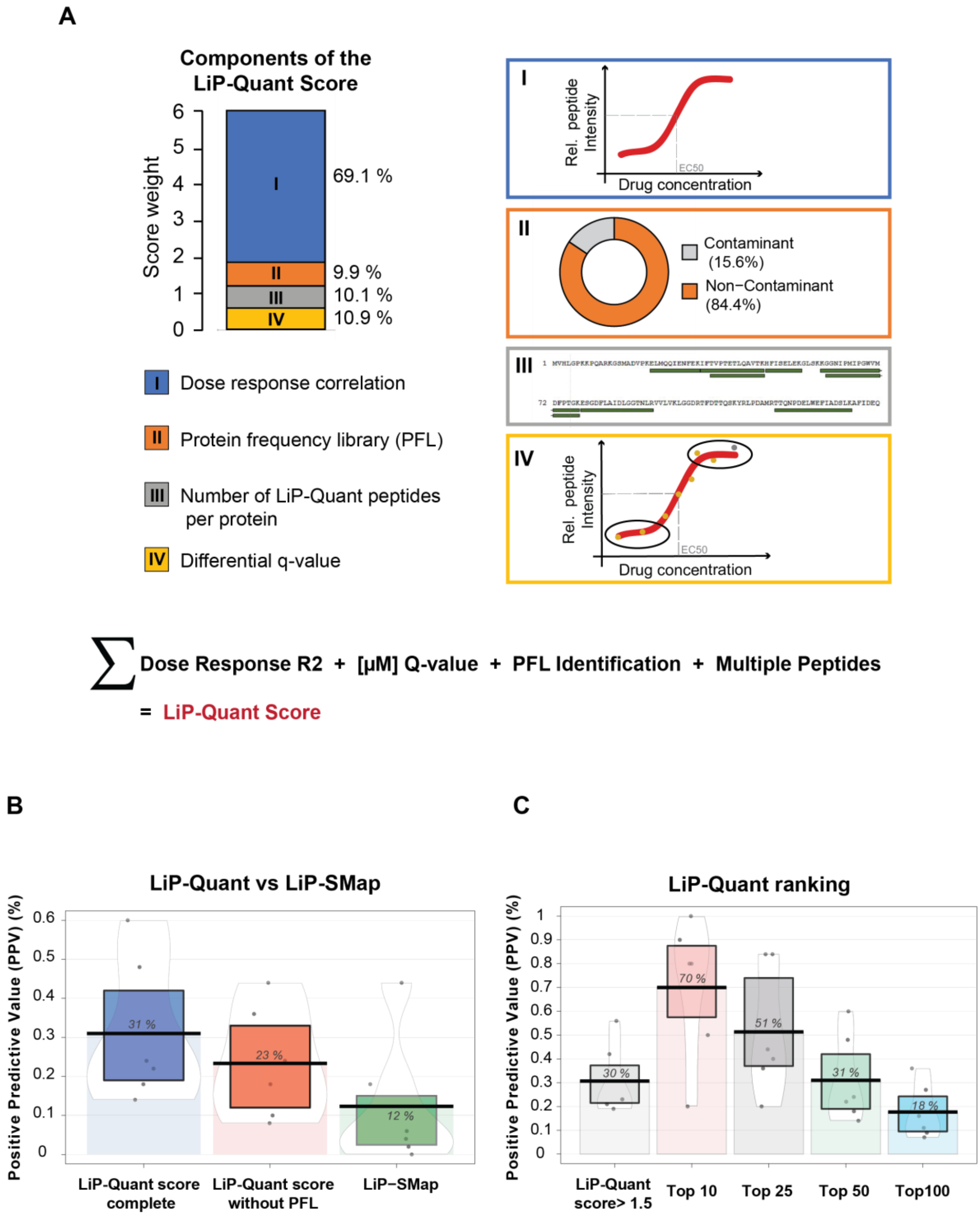


**FIGURE S1: Development and verification of the automated LiP-Quant analysis pipeline: 1A:** Ranking by LiP-Quant score is based on 4 components including the correlation coefficient to a sigmoidal trend of the isothermal dose-response profile (I), the protein's frequency as a common contaminant in LiP-Quant experiments (Protein Frequency Library, PFL) (II), the number of LiP-Quant peptides assigned per protein (III) and the statistical significance of the relative peptide abundance between two points (IV). The stacked histogram shows the machine learning-derived relative weight of each component expressed as percentage of the LiP-Quant score over a possible maximum score of 6. **1B:** Positive Predictive Value or precision of 3 different protein-small molecule interaction predictors: LiP-Quant using all 4 components of the LiP-Quant score (see **S1A**), LiP-Quant excluding the common contaminant filtering by PFL (see **S1A**) and LiP-SMap. The boxes show the interval of the interquartile range and the bean plots the smoothed density curve relative to the full data distributions. The horizontal bars represent the means of the number of true positives peptides found in the experiments shown in this work. **1C:** Positive predictive value when considering as positive hits the peptides with LiP-Quant score > 1.5, or the top 10, 25, 50 and 100 ranking peptides. Peptides scoring > 1.5 are generally contained in the top 50 scoring peptides. The positive predictive value and data representation calculations are the same as in 1B.

A

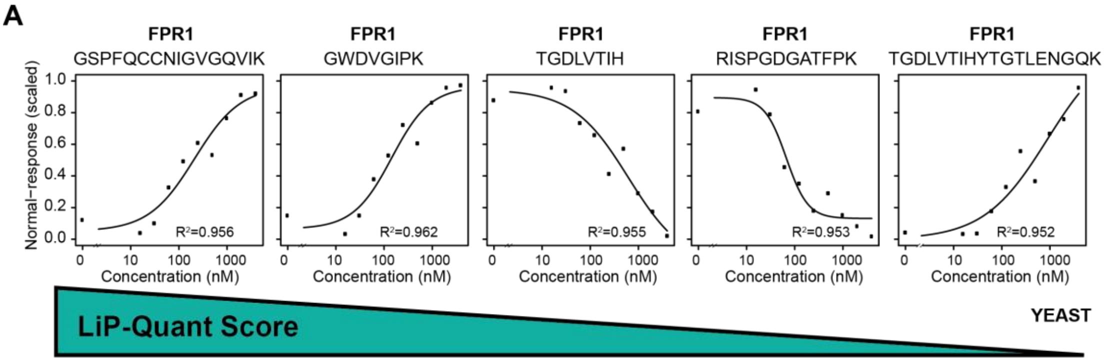


B

**FPR1**

| LiP-Quant peptide | $R^2$ | Score | Rank |
|---|---|---|---|
| **GSPFQCCNIGVGQVIK** | 0.956 | 3.228 | **1** |
| GWDVGIPK | 0.962 | 3.111 | 2 |
| TGDLVTIH | 0.955 | 3.070 | 3 |
| RISPGDGATFPK | 0.953 | 2.974 | 4 |
| TGDLVTIHYTGTLENGQK | 0.952 | 2.771 | 5 |

C

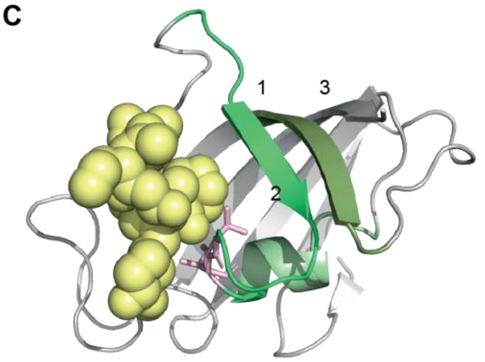


D

**Other protein targets**

| LiP-quant peptide (protein) | $R^2$ | Score | Rank |
|---|---|---|---|
| AHLVAIEKPECTGQR (**ARI1**) | 0.969 | 2.711 | 6 |
| GDLVITEESWNK (**ARI1**) | 0.952 | 2.581 | 8 |
| APIVAYAELIAAR (**SYEC**) | 0.937 | 2.510 | 10 |

E

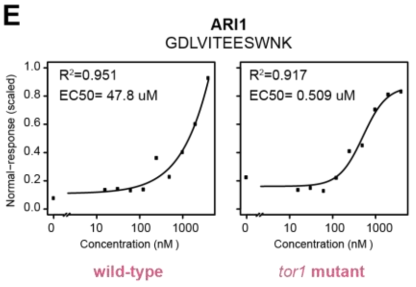


F

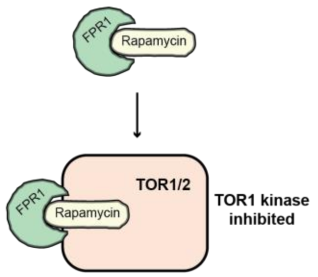


G

| Stripped Sequence | Directionality | Score | LiP Rank | Compound | Log2 Fold Change |
|---|---|---|---|---|---|
| GWEEGVAQMSVGQR | Increasing | 4.91 | 1 | Rapamycin | 2.754 |
| | | 3.82 | 1 | FK506 | 4.046 |
| RGQTCVVHYTGMLEDGK | | 4.49 | 2 | Rapamycin | 2.715 |
| | | 3.49 | 3 | FK506 | 4.679 |
| LTISPDYAYGAT | Decreasing | 3.10 | 5 | Rapamycin | -2.130 |
| | | 1.72 | 11 | FK506 | -0.730 |
| YTGMLEDGK | | 3.07 | 7 | Rapamycin | -2.250 |
| | | 3.01 | 4 | FK506 | -1.270 |
| LVFDVELLK | | 3.01 | 8 | Rapamycin | -2.145 |
| | | 1.86 | 8 | FK506 | -1.489 |
| VFDVELLK | | 2.35 | 13 | Rapamycin | -2.122 |
| | | 1.73 | 10 | FK506 | -0.969 |

**FIGURE S2: Benchmarking LiP-Quant in yeast cells and consistency of LiP-Quant with drugs targeting the same binding sites: S2A:** Dose-response curves showing relative intensities of LiP-Quant peptides obtained from *S. cerevisiae* lysates over a rapamycin concentration range (0 nM to 2000 nM). Curves of the top 5 LiP-Quant peptides ranked by LiP-Quant score are shown, all of which are from FPR1, the expected direct target for this drug in yeast cells. **S2B:** LiP-Quant peptides ranking in positions 1-5 of the LiP-Quant experiment done with rapamycin in *S.cerevisiae* cells. All 5 are FPR1 peptides. **S2C:** Structural model of the holocomplex of FPR1 with the rapamycin analog FK506 (PDBid: 1yat). The green electron density corresponds to the top 3 FPR1 LiP peptides and the drug ligand is shown in yellow. Valine and isoleucine residues in direct contact with the compound are shown with pink sticks. The 3 top scoring peptides are positioned within the binding site of rapamycin. **S2D:** LiP-Quant peptides ranking in positions 6, 8, 10 of the LiP-Quant experiment done with rapamycin in *S.cerevisiae* cells. **S2E:** Dose-response curves of the relative peptide intensity of GDLVITEESWNK, which maps to ARI1, over a concentration range of rapamycin lysates of wild type or *tor1-fpr1* mutant yeast. **S2F:** Model of TOR1 kinase inactivation through the binding of rapamycin to FPR1. **S1G:** LiP-Quant peptides generated with LiP-Quant are consistent across drugs that bind the same site, as the same specific peptide sequences were generated in both the rapamycin and FK506 assays. In both experiments the LiP-Quant peptides show similar regulation, with peptide intensity changes in the same direction (log2 fold change) upon drug binding.

**A**

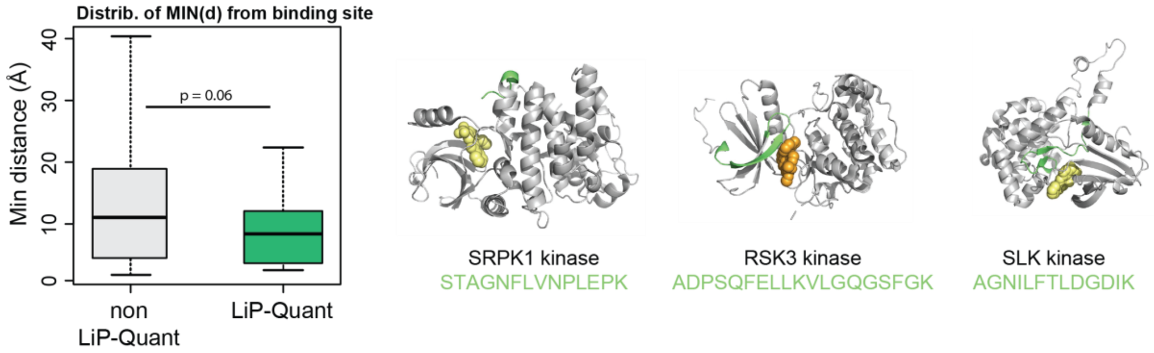


**B**

**NQO2**

| LiP-Quant peptide | $R^2$ | EC50 | Rank | Compound |
|---|---|---|---|---|
| EEPIPCTAHWHFGQ | 0.967 | 13662.83 | 2 | Selumetnib |
| EEPIPCTAHWHFGQ | 0.912 | 2600.00 | 8 | Staurosporine |
| DITGTLSNPEVFNYGVETHEAYK | 0.873 | 926.29 | 14 | Staurosporine |

**C**

**PP2A**

| LiP-Quant peptide | R2 | EC50 | Rank |
|---|---|---|---|
| YSFLQFDPAPR | 0.990 | 18.22 | 1 |
| NVVTIFSAPNYCYR | 0.984 | 19.28 | 2 |
| AHQLVMEGYNWCHDR | 0.979 | 11.38 | 3 |

Median EC50
18.23 nM

**PP1A**

| LiP-Quant peptide | R2 | EC50 | Rank |
|---|---|---|---|
| IKYPENFFLLR | 0.951 | 663.0 | 4 |
| EIFLSQPILLELEAPLK | 0.946 | 55.47 | 5 |
| IYGFYDECKR | 0.953 | 63.26 | 7 |

Median EC50
63.26 nM

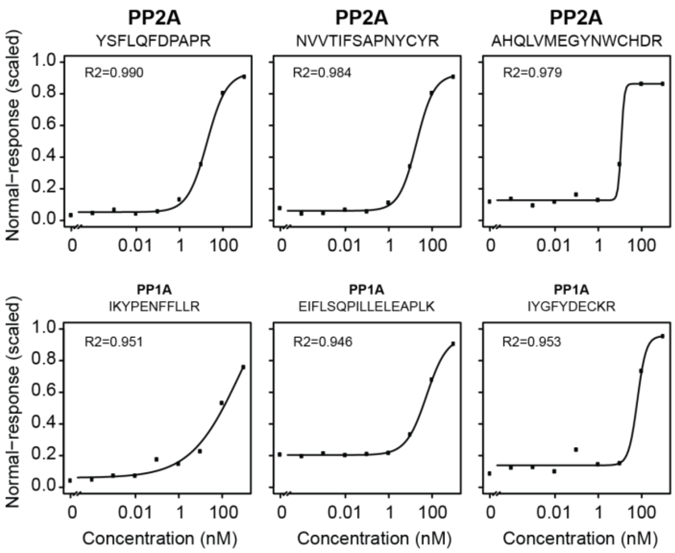


**D**

| Stripped Sequence | Directionality | Score | LiP Rank | Compound | Log2 Fold Change |
|---|---|---|---|---|---|
| NVVTIFSAPNYCYR | Increasing | 4.82 | 2 | Calyculin A | 1.223 |
| | | 2.76 | 2 | Fostriecin | 0.838 |
| AHQLVMEGYNWCHDR | | 4.61 | 3 | Calyculin A | 0.992 |
| | | 2.59 | 3 | Fostriecin | 0.636 |
| GAGYTFGQDISETFNHANGLTLVSR | | 3.62 | 8 | Calyculin A | 1.510 |
| | | 1.61 | 21 | Fostriecin | 0.884 |
| SPDTNYLFMGDYVDR | | 3.35 | 10 | Calyculin A | 0.730 |
| | | 1.56 | 30 | Fostriecin | 0.466 |

**FIGURE S3: Structural and quantitative analysis of two classes of druggable targets: kinases and phosphatase: S3A**: Distribution of minimal distances from the drug binding site calculated for LiP and non-LiP peptides identified by the LiP-Quant experiment with staurosporine. Holocomplexes of kinases bound to the kinase inhibitors or ATP were used for this analysis. The median distance from the drug binding site is 8.3 Å for LiP-Quant peptides and 11.4 Å for non-LiP-Quant peptides. Representative holocomplex structures of SRPK1, RSK3 and SLK kinases with their drug or ATP ligands are shown. LiP-Quant peptides are colored in green; drug ligand in yellow, ATP in orange. **S3B:** LiP-Quant peptides mapping to a candidate off-target protein of selumetinib and staurosporine. **S3C:** Top 6 ranking LiP-Quant peptides in the LiP-Quant experiment with calyculin A; these peptides map to PP2A and PP1A. LiP-Quant peptide intensities in response to drug dose are also shown. **S2D**: LiP-Quant peptides generated with LiP-Quant reflect a consistent binding mechanism at the same site of PP2A, as the same specific peptide sequences were generated in both the fostriecin and calyculin A assays. In both instances the LiP-Quant peptides show similar regulation with peptide intensity changes in the same direction (log2 fold change) upon drug binding.

A

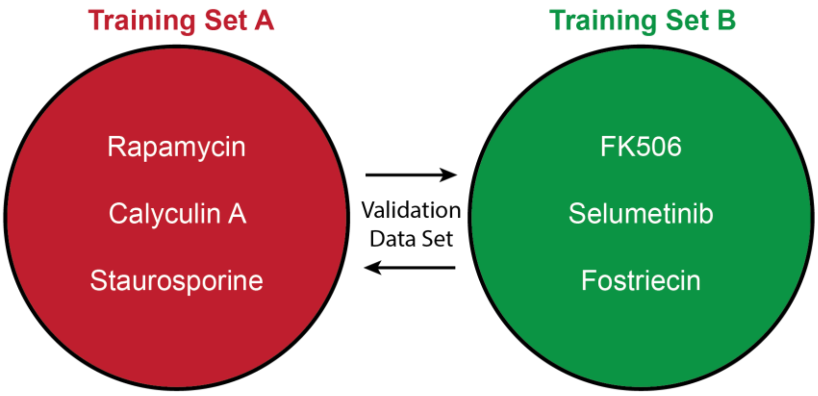


B

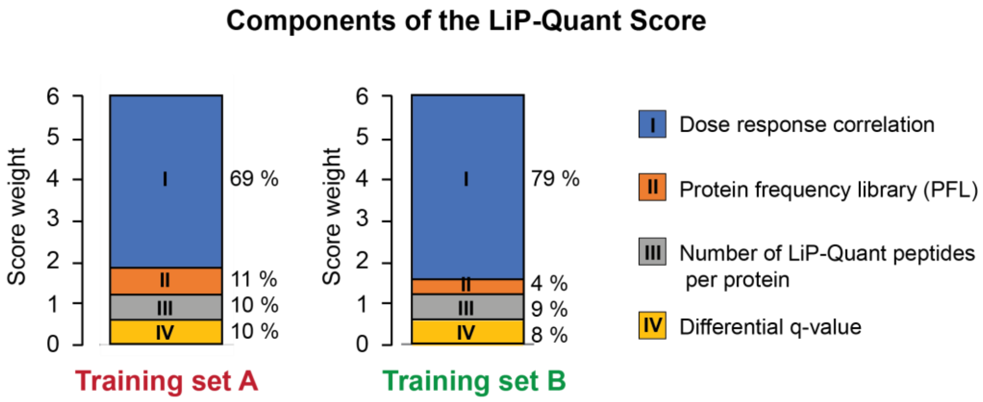


C

| | | | *LiP-Quant Score* | | *LiP Rank* | |
|---|---|---|---|---|---|---|
| **Compound** | **Sequence** | **Genes** | **Set A** | **Set B** | **Set A** | **Set B** |
| Rapamycin | GWEEGVAQMSVGQR | FKBP1A | 4.910 | 4.940 | 1 | 1 |
| | RGQTCVVHYTGMLEDGK | FKBP1A | 4.490 | 4.492 | 2 | 2 |
| | RGQTCVVH | FKBP1A | 3.775 | 3.786 | 3 | 3 |
| | VFDVELLKLE | FKBP1A | 3.338 | 3.344 | 4 | 4 |
| | LTISPDYAYGAT | FKBP1A | 3.100 | 3.044 | 5 | 6 |
| | EEDEEEDVPGQAKDEL | CALR | 3.069 | 3.125 | 6 | 5 |
| | YTGMLEDGK | FKBP1A | 3.067 | 3.005 | 7 | 7 |
| | LVFDVELLK | FKBP1A | 3.014 | 2.890 | 8 | 8 |
| | LVFDVELLKLE | FKBP1A | 2.893 | 2.769 | 9 | 10 |
| | DYPGLPVFLLGH | MGLL | 2.875 | 2.808 | 10 | 9 |
| FK506 | GWEEGVAQMSVGQR | FKBP1A | 3.818 | 3.852 | 1 | 1 |
| | GVQVETISPGDGR | FKBP1A | 3.726 | 3.702 | 2 | 2 |
| | RGQTCVVHYTGMLEDGK | FKBP1A | 3.488 | 3.580 | 3 | 3 |
| | YTGMLEDGK | FKBP1A | 3.007 | 2.976 | 4 | 4 |
| | TISPGDGR | FKBP1A | 2.813 | 2.755 | 5 | 5 |
| | EEDEEEDVPGQAKDEL | CALR | 1.954 | 1.853 | 6 | 6 |
| | GWEEGVAQ | FKBP1A | 1.939 | 1.790 | 7 | 7 |
| | LVFDVELLK | FKBP1A | 1.856 | 1.716 | 8 | 9 |
| | EFIEGVSQFSVK | PPP3R1 | 1.851 | 1.731 | 9 | 8 |
| | VFDVELLK | FKBP1A | 1.731 | 1.641 | 10 | 10 |

**FIGURE S4: LiP-Quant pipeline development, training and validation: S4A:** The schematic illustrates the ground truth experiments used to determine sub-score weightings for defining the LiP-Quant score. Two independent data sets, each from three experiments using three drugs with known targets, were analyzed by linear discriminant analysis to determine parameters that contribute to positive target identification. The weightings derived from each of these training sets was then validated on an independent data set, again each consisting of three experiments using three different drugs with known targets (i.e. trained on A and tested on B and vice versa). **S4B:** Four parameters were identified that contribute to positive target identification for each training set. The criteria include the correlation coefficient ($R^2$) to a sigmoidal trend of the isothermal drug dose-response profile (69% of the LiP score), whether the identified protein is a common contaminant in LiP-Quant experiments (Protein Frequency Library (PFL), 10% of the LiP score). Extra parameters are the number of LiP peptides per protein and the statistical significance of the relative abundance of a LiP peptide between drug and vehicle treated samples. The weightings of these parameters were found to be very stable. **S4C:** LiP-Quant peptides identified, scored and ranked are consistent when analyzed with weightings from either training set. Shown are LiP-Quant scores and ranks for one experiment from training set A (rapamycin) and from training set B (FK506) in two experiments.
